# A universal metric for evaluating, optimising and benchmarking the performance of a Research Technology Platform (RTP)

**DOI:** 10.1101/133207

**Authors:** A.M. Petrunkina, A. Filby

**Affiliations:** Department of Medicine, University of Cambridge School of Clinical Medicine, Box 157, Addenbrooke’s Hospital, United Kingdom; Faculty of Medical Sciences, Newcastle University, Framlington Place, Newcastle upon Tyne, NE1 7HH, UK

**Keywords:** operations, metrics, performance, optimisation, Science/Research Technology Platform

## Abstract

Research Technology Platforms (RTPs) exist to facilitate the application and utilisation of specific analytical technologies to the highest possible standard thus delivering reputable data across a broad spectrum of research themes. Specifically, RTPs centralise expertise in a given technology and provide an unparalleled level of continuity and practical knowledge retention that simply cannot be achieved by more organic, ad hoc means of support. As small non profit businesses often tasked with recovering all or a percentage of their running costs, RTPs are under significant pressure to keep pace with rapidly advancing technology and new methodologies against a back drop of dwindling funding for scientific research. At present there are a number of non-trivial issues that make assessing the operational performance of a RTP difficult to determine on a standalone basis let alone attempting to benchmark against other RTPs within the same or different technology fields. Firstly, depending on the technological speciality the RTP may work to one of essentially three operational models. RTPs such as Bio-Imaging or Cytometry provide access to well-maintained analytical systems that can be utilised by trained individuals for a timed access charge. In some cases there will be a requirement for assisted operation of certain instruments by core staff (e.g. cell sorters). Genomics and Proteomics RTPs tend to function on a project basis whereby users will not access the technology themselves rather pay for a full analytical service often with a milestone-based approach for tracking progress. Other RTPs work to a hybrid approach were technical staff provide certain elements of sample preparation for specific projects prior to analysis on core supported, user accessible instrumentation. Secondly the specific operational costs that each RTP is tasked to recover varies significantly on a local, national and international level due to institutional subsidies. These operational costs can include staff salaries, instrument maintenance, associated running consumables, and in some cases instrument depreciation but there is standardised rule as to what each RTP is tasked to recover and to what percentage.

Here we present a generalised mathematical approach to describe the customisable metrics of any given RTP serviceThe general strategy how to increase performance within the framework of this approach has been identified through breaking down these customisable metrics into components and maximising them according to specific requirements. These strategies could be potentially adopted for different operational or local procedures, integrating the specifics related to the institutional or national policies. The approach laid down here should be considered as a trigger for opening a discussion around how to address optimising RTP performance and allow for benchmarking across the full breadth of RTPs.

## Introduction

Many RTPs are faced with the increased demand in usage which may potentially represent a major operational issue, especially in the infrastructure-lacking environment and in context of competing for limited funding (Haley, 2009).

Strategies to optimize operations and management of such research laboratories, operating as small businesses for the purpose of provision of the specialist expertise and equipment, require identification of carefully selected metrics for evaluating general operations and management (Turpen et al, 2016). In particular, they would require adopting a generalist approach that will be adaptable to and compatible with the local specifics and policies and be universal, e.g. could be applied to variety of services. Such an approach utilising as a tool a mathematical algorithm compatible with the local specifics and policies would be instrumental in increasing performance of RTP as a successful enterprise. The limiting factor so far remains the commonly widespread misapprehension that no generalised strategy is possible for quantifying the capacity and utilization of services and efficiency of operations per se. That is because the specialist requirements (e.g. requirement for a dedicated operator or different requirements of a service) are too specific for the core services of different nature to establish and apply uniform/universal metrics across the broad range of services and because any particular approach or idea adopted in one country how to measure the capacity and output may collide with institutional or country-specific policies elsewhere and thereby prevent translational applications across the institutions and borders. Feedback received with respect to our previous work describing the algorithm for estimating capacity of the specialist sorting service (Petrunkina, 2013) has highlighted two major vulnerabilitites of the proposed non-generalist solution: on one hand, its specificity for the sorting operator service perceived as the lack of the generalised approach that would be applicable to any science technology platform and any service, and, on another hand, its potential for misinterpretation as a tool to predict capacity accurately.

RTPs are often judged against two performance criteria; cost recovery and links to scientific output. There are several issues with this however. Firstly RTPs are not “masters of their own destinies” and often rely on academic users securing grants to fund research and related usage of a given set of technologies. While the presence of a well-run, highly competent RTPs may help researchers demonstrate that they have the technological backing to deliver on a research idea, it is the idea itself and the reputation of the researcher that will ultimately dictate the success or failure of a grant application. In a similar vein, it is not appropriate to measure the performance of a RTP on scientific output (publications etc.) unless the RTP itself is officially tasked with publishing work.

Here we present a standardised mathematical, yet highly practical and accessible method for assessing the operational performance of any RTP. Moreover our approach provides a justifiable method to calculate the practical capacity of any aspect of a RTP service for the purpose of calculating re-charge rates as function of practical capacity.

The results of our method can be used to evaluate all aspects of the service from staffing levels, activities and gives an all important measure of efficiency. Science and technology are fast moving fields and it is essential that RTPs are able to identify areas of weakness and thus opportunity for improvement and growth so as to adapt quickly to changes/needs.

## Tracking capacity and efficiency of RTP - practical example of calculation

Let us consider a RTP providing service in sample processing and analysis (e.g. Phenomics, Genomics or Proteomics). The major services delivered by this unit can be broken into following 6 components: provision of sample service, training, maintenance, administration, research, staff management. The RTPRTP is staffed by team of three (3 FTE): core manager, specialist and assistant, with varying job descriptions as summarised in the Table 1. Under local labour legislation (United Kingdom in this example) and local institutional policies (academia), we have to account for 8 public holidays, 253 working days, annual leave entitlement for academic staff of 33 days, annual leave entitlement for technical staff of 28 days, contractual working hours for academic staff – minimum of 38.5 hours per week, contractual working hours for technical staff - on average 36.5 hours per week. We could summarise all this information in the Table 2, the cumulative working times being of 4978 hrs p.a. Let us assume that the number of instruments used for sample service is two (*k*_*ss*_ =2), and the non-operative local preparation time (non-bookable) for each systemn is 1.5 hrs (45 min, cleaning and 45 min shutdown), resulting in 3 hrs on non-operative time a day, or cumulatively, 253·3=759 hrs per annum. Apart from providing sample preparation and operator service on these two instruments, the RTP also has three user-operation instruments (*k*_*so*_=3).

**Table 1.**
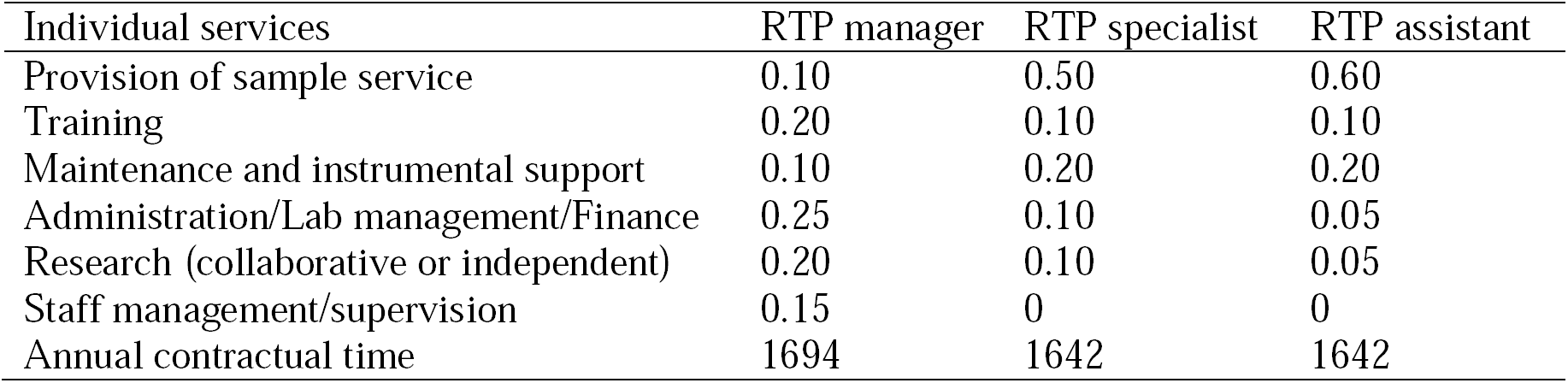
Matrix P for RTP providing sample service

**Table 2:** Cumulative service times for each component of services.

| Individual services | □ for all staff members | Hours, p.a. |
| --- | --- | --- |
| Provision of sample service | $0.10 \cdot 1694 + 0.5 \cdot 1642 + 0.6 \cdot 1642$ | 1976 |
| Training | $0.20 \cdot 1694 + 0.1 \cdot 1642 + 0.1 \cdot 1642$ | 667 |
| Maintenance and instrumental support | $0.10 \cdot 1694 + 0.2 \cdot 1642 + 0.2 \cdot 1642$ | 826 |
| Administration/Lab management/Finance | $0.25 \cdot 1694 + 0.1 \cdot 1642 + 0.05 \cdot 1642$ | 670 |
| Research (collaborative or independent) | $0.20 \cdot 1694 + 0.1 \cdot 1642 + 0.05 \cdot 1642$ | 585 |
| Staff management/supervision | $0.15 \cdot 1694$ | 254 |
| Instrument availability | $1642 \cdot K - M - T$<br>K-number of instruments, M-maintenance, T- training | For capacity and billing purposes |

Following service outputs have been recorded:

Sample service delivered during year in question was 1000 hrs. Closures were for 6 days (Christmas week and refurbishments).

Authorised absences: 36 days for the manager (33 days leave and 3 days conference), 33 days for the specialist (28 days leave and 5 days training course), 30 days for the assistant (28 days leave and 2 days sickness)

Breakdown recorded for instruments with operator service: 1 day instrument A, 3 days instrument B, no breakdowns for user-operated instruments. Annual levels of maintenance were 600 hours. Training delivered on self-operation instruments: 300 hrs. 4800 hrs usage on end-user operated equipment were recorded.

However, how can one translate these operational outputs into standardised metrics of utilization and efficiency applicable to other similar resources?

We could calculate the cumulative time available for all services as the sum of time dedicated by each member of RTP to the service provision less non-operative time (e.g. time used for the preparation of machinery of work place). According to the Table 2 and equations [6.1-6.3] from the Appendix, the theoretical capacity for sample service amounts to 1976 hrs, less non-operational time of 759 hrs yields 1217 hrs. Forthwith, the theoretical capacity per operator per day will be obtained by dividing the theoretical capacity by the number of working days and the number of operators of service (253x3 in this case): 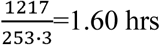.

We can now calculate operational days (OD) and efficiency coefficient (E) taking into account data from above for closures (6 days), cumulative authorised absences (99 days) and breakdowns (1/2 and 3/2 days):

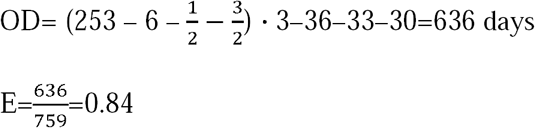

Practical capacity=1217 hrs ·0.84=1020 hrs

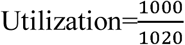.100%= 0.99. 100%, or 98% of the available staff capacity.

Theoretical instrumental capacity for instruments C, D and E (self-service) = 1642·3= 4926 hrs according to [8.1] and the practical instrumental capacity for self-operation instruments is 1642·3-600-300=4026 hrs according to [8.2] from Appendix 2.

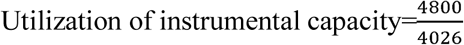

The usage levels for operator sample service are close to 100% and the efficiency is high (83%), so no immediate action is required. However, for the user-operated instruments, the recorded levels are above and beyond capacity based on normal working times, thus a review of operational strategy for sample service delivery to take off the load from self-operation service might be considered. However, a much more straightforward would be to consider a purchase of a new instrument for self-service because the transition from self-service into operator service is usually less expedient and not cost-effective (while a vice-versa transition should always be considered as an option if proven technically feasible, viable and reasonably practicable).

## Tracking capacity and efficiency of a RTP - a generalised mathematical approach

Based on the practical example above, we can introduce mathematical definitions and equations to define metrics of performance for a shared laboratory resource, staffed with *n* members of staff and providing *m* different specialist services across the portfolio range. In particular, practical capacity, utilization of practical capacity and efficiency will be defined based on staff roles descriptions, operational strategy, service records and number of operational days when service was provided. For any resource, these parameters can be calculated retrospectively using the working spreadsheet (supplementary data) and widely used for evaluation, strategic and operational planning. Further, historical data can be used for maximising the capacity and efficiency and optimisation of services as described below.

## Maximising the capacity and efficiency of a RTP – operational applications of the approach

Any service directly associated to the requirement of a dedicated operator usually implies that only a rather narrow spectre of solutions is available to increase this capacity. Strategies are most often reduced to buying a new instrument, implying an increase in the number of FTEs required to operate it, or most likely increasing dramatically the workload of the current staff, all of which will inevitably have a negative impact on quality or RTP performance. A practical example associated with the provision of cell sorting service, and a recent algorithm by Petrunkina (2013) have captured the metrics of capacity and efficiency as applied to this particular service. With the advent of the new and more automated cell sorters, different and more creative solutions to increase cell sorting capacity could be implemented, including user-operated self-service sorting. However, it is possible to present additional approaches to optimize usage of the current instrumentation, independent of their level of automation. The improvements may be those of technical nature (such as small technical implementations to reduce instrument setup), of outreach and educational nature (e.g. more comprehensive training for advanced users and provision of instrument troubleshooting guides), or of administrative and managerial nature (booking strategies to optimize resource usage, staggering shifts within extended hours or introducing late shifts to supervise and assist with experiments that cannot be scheduled within normal working hours, e.g. for clinical reasons). The common consensus is that it is not possible to provide the universal guidance across all specialist RTPs, although this default agreement may be based on a fallacy. While one definitely has to agree that there no identical step by step course of actions that will fit all facilities as some of these approaches may collide with institutional or country-specific policies, it is possible to present a generalised model which can be broken into components, each of which can be addressed separately according to the specifics of the case. In particular, optimisation strategies for improving the service and maximising performance will be constrained by the specifics of the service, local, institutional or national policies and legislation.

As one can see from the above example, no specific assumptions have been made about the nature of services and operations, instrumentations or other restrictions. All these details as well as the details related to the country-specific labour legislation (i.e. UK in our example) or to the institutional policy can be easily identified, accounted for and substituted. This stepwise procedure gives a very clear illustration that the above mathematical approach can be applied to any service or RTPas long as the facility is prepared to undergo an operational review and optimise their task allocation/operational strategies (see Box et al. 2012). It implies that this approach could be applied to phenotyping, histology, high content, cytometry, imaging and many other facilities and science technology platforms (see flow chart in the supplementary Figure 1). It must be emphasised that this approach is not an attempt to predict the practical usage for any service which would not been possible given the variety of contributing factors (instrument durability, human factor including skills, competencies and behaviour and other unforeseen circumstances). Much more, this approach is focussed on regular monitoring of the utilization of the practically available capacity (which is being evaluate retrospectively and will always differ from theoretical predictions) and efficiency of RTP and immediate implementation of the optimisation strategy based on the analysis of these metrics and contributing factors.

Indeed, everything required for working out the robust optimisation strategy is available in the equations [6.2], [6.3] and [7.1] from the Appendix and the optimisation flow chart. To increase practical capacity one needs to maximise the operative time and efficiency, or, in other words, maximise the difference between cumulative time assigned to each service t and non-operative time d [vector (t-d)] and to maximise the number of operational days OD. According Eq. 6.2, one can achieve that by maximising time *t* dedicated to a particular service (through employing new FTE, revising operational strategy, updating job descriptions and implementing multi-tasking) and by minimising non-operative time *d* (through using automated instrumentation, modifying existing instrumentation, reviewing SOPs and being innovative). Indeed, the aspects of incorporating multi-tasking or parallel operation, giving the users access to self-operation will increase nominal operative time. Equally, reviewing or updating job descriptions according to the dynamic operational strategy will allow re-populating matrix P by re-assigning the tasks/ times dedicated to each particular service. If automated instrumentation can be used, and latest technical solutions can be implemented, then less preparation and calibration time will be required, which will decrease non-operative time and increase operative time. Finally, such a simple aspect as a comprehensive user consultation, booking questionnaires to pre-qualify what is required and advanced thinking to plan the optimal configuration will maximise use of the available operative time and allow to run more cells/samples through the instrument within given time. Increasing the number of operational days will boost the efficiency (Eq. 6.3) – that can be achieved by keeping the downtime low, planning for upgrades and preventive maintenance visits to be done on the same day, maintain the right balance of the professional and personal development in order to enhance staff satisfaction (to avoid understaffing due to turnover or sick days) but without endangering the capacity of service. All that will allow booking more time for the service according to the increased practical capacity, and, thereby, increase utilization rates (Eq. 7.2).

Apart from the applications associated with assessing the output, metrics and benchmarking of service, this algorithm could be also used for accounting, budgeting, setting the user charges and supporting funding applications for staff and equipment. The practical staff and instrumental capacity estimates based on prior analysis can be used for forecasting estimates and thus have great potential to become universal operational metrics for varied aspects of RTPenterprise.

## Maximising the capacity and efficiency – practical examples

Let us consider a shared laboratory resource from the previous example providing service in sample processing and analysis (e.g. phenotyping, genotyping or proteomics). All assumptions about the job descriptions and legislation are the same, however, in the first scenario the service output is supposed to be significantly lower, and in the second scenario the waiting times for samplesprocessingarelongerthan4weeks.

### Hypothetical situation A: underutilization of resources

Following service outputs have been recorded:

Sample service delivered during year in question was 500 hrs. Closures were for 19 days (X-mas, Easter week and delayed refurbishments). Authorised absences: 53 days manager (33 days leave, 10 days conference, 10 days consultancy), 38 days specialist (28 days leave and 10 days training course), 40 days assistant (28 days leave and 12 days sickness).

Breakdown recorded for instruments with operator service: 5 days instrument A, 10 days instrument B, no breakdowns for user-operated instruments.

According to [6.1-3] and Table 2, the theoretical capacity for sample service amounts to 1976 hrs, less non-operational time of 759 hrs yields 1217 hrs.

Capacity per operator per day 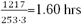

We can calculate operational days taking into account data from above for closures (19 days), cumulative authorised absences (131 days) and breakdowns (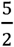 and 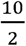days):

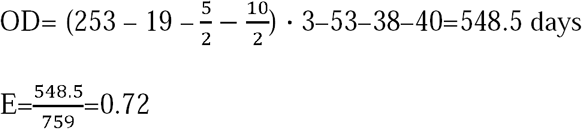

Practical capacity=1217 · 0.72=876 hrs

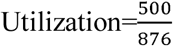 · 100% 0.57 or 57% of available capacity

Both the capacity and the utilization levels are very low – less than 60% of the available practical capacity for the operator-delivered service is utilized. A thorough review of operational objectives for sample service delivery and their tactical execution is overdue. The aspects of marketing the service (to increase low usage), outreach and educational activities should be considered to attract more users for the service.

On another hand, the efficiency is considerably lower than benchmark of 80%. To increase efficiency, one needs to increase numbers of operational days. Simultaneously, one needs to look into reasons for the low efficiency, in this case associated with prolonged authorised absences and breakdowns. In this scenario, it was immediately apparent that staff absence levels and breakdowns are very high, possibly due to other shortcomings in tactical execution (e.g. inadequate prioritization of tasks or lack of leadership, poor practices), although attention needs to be paid to the high sickness levels whether they may have be due to occupational reasons. In this particular situation, there is no need to purchase additional instruments or employ new staff to increase capacity because the optimisation potential of the existing instrumental and staff capacity is by far not exhausted.

### Hypothetical situation B: shortage of instrumental and/or staff practical capacity – multiple-solution approach

In this case we have the same conditions as described above in the section ‘Tracking capacity and efficiency of shared laboratory resources - practical example of calculation’ practical operational capacity of 1010 hrs a year for the operator-delivered service but except that there are long waiting times for the service, indicating that the demand overstrips the practical capacity. Because the efficiency is above the benchmark for a well-run resource (E=0.84), and the utilization is nearly 100% (=98%), one should consider several avenues for increasing the practical capacity that are not related to raising awareness of technology or facility marketing, and are addressing the operational strategy and operational processes themselves rather than straightforward optimisation of the number of operational days (minimising absences and breakdowns). Those avenues could include but not be limited to 1) decreasing demand for operator service by attempting expanding self-operation, 2) increasing practical capacity through introducing new SOPs involving multi-tasking, such as coordinated preparation of two units in the mornings to reduce non-operative time, 3) re-allocation of tasks from areas where offer outstrips the demand, et cetera. It could be realised by re-writing matrix P (job descriptions) to ensure more time is available for operator-delivered sample service, if that would not affect negatively other aspects of operations.

A consideration should be given to enhancing outreach and training in order to increase the levels of self-operated service (that may provide relief for the oversubscribed operator sample service). According to the Table 2 (product of Matrix P of duties and working times for each member of staff), staff capacity for provision of training is 667 hrs. Only 300 hrs of training was delivered. Therefore, there is training potential available. However, instrumental capacity for self-operation was already fully exhausted (119%). Therefore, if the manager would favour the promoting self-operation, they would need to apply for funding for a new instrument which could boost the self-operation, relieve the exhausted practical capacity for operator driven service and ensure there is a solid back up if one of the instruments is down. Under existing conditions training new users alone without expanding equipment base would only add to frustration because no instrumental capacity is available.

If no funding is available or the procurement can only happen in the mid-term, the manager shall consider updating job descriptions to increase time allocated to the operator service, *t*_*i*_. From the service output record and the Table 2 it is apparent that at least 300 hrs are under-delivered for training. These ‘free’ hours could be re-assigned to the provision of sample service. Moreover, the maintenance demand seems to be overestimated – according to the current job descriptions maintenance amounts to 826 hrs, but with only 600 hrs delivered, the current equipment has marginally no downtime. Clearly, some fraction of time could be reassigned to sample service (while some surplus provisions must be must be left both for training and maintenance to enable quick response in emergencies and to allow for the maintenance on additional equipment shall that be purchased).

Beyond that, after reviewing procedures across peers (survey of national facilities), the manager has detected that the preparative non-operational time is fairly high – 1.5 hrs a day (45 min each kit) while other sites report in average 30 min per kit. Practical capacity can be increased by minimising non-operative times d_*i*_. Unless there a specific operational reason why both kits cannot be started up by the same operator in parallel (multi-tasking) in this particular location (there may be a unique explanation, for example handing different microorganisms requiring specific SOPs), a standardised solution would be more efficient. For example, re-write the start-up SOP accommodating a parallel start-up of units and QA.

Such re-distribution of duties following thorough review and new organisational approach would result in a modified allocation of working times to services and their cumulative working times. Moreover, it will affect the calculation of practical capacity. Indeed, the Table 3 (optimised Table 1 lists the re-organised distribution of duties for RTP specialist and RTP assistant who now provide less training (5% instead of 10% each for each service freeing up additional 20% of FTE) but more sample service (60% and 70 % instead of 50% and 60%). Table 4 (optimised Table 2) illustrates the new breakdown of the cumulative service times of 4978 hrs. Through changing SOP for start-up, the non-operative service time was reduced to 1 hr instead of 1.5 hrs (two kits in parallel take 1 hrs instead of 2x45 min). Therefore, daily non-operative time is now 2.5 hrs instead of 3 hrs, and the annual non-operative time is 253x2.5= 632.5 hrs instead of 759 hrs. In total, the available sample service booking time (practical capacity) is now 2304-632.5= 1671.5 hrs (instead of 1217 hrs). That is 37% increase in practical capacity and will help to reduce long waiting time without involving hiring new staff or introducing long working hours.

**Table 3.** Optimised Matrix P for RTP providing sample service

| Individual services | RTP manager | RTP specialist | RTP assistant |
| --- | --- | --- | --- |
| Provision of sample service | 0.10 | 0.60 | 0.70 |
| Training | 0.20 | 0.05 | 0.05 |
| Maintenance and instrumental support | 0.10 | 0.15 | 0.15 |
| Administration/Lab management/Finance | 0.25 | 0.10 | 0.05 |
| Research (collaborative or independent) | 0.20 | 0.10 | 0.05 |
| Staff management/supervision | 0.15 | 0 | 0 |
| Annual contractual time | 1694 | 1642 | 1642 |

**Table 4:** Cumulative service times according the optimised matrix P for service delivery

| Individual services | □ for all staff members | Hours, p.a. |
| --- | --- | --- |
| Provision of sample service | $0.10 \cdot 1694 + 0.6 \cdot 1642 + 0.7 \cdot 1642$ | 2304 |
| Training | $0.20 \cdot 1694 + 0.05 \cdot 1642 + 0.05 \cdot 1642$ | 503 |
| Maintenance and instrumental support | $0.10 \cdot 1694 + 0.15 \cdot 1642 + 0.15 \cdot 1642$ | 662 |
| Administration/Lab management/Finance | $0.25 \cdot 1694 + 0.1 \cdot 1642 + 0.05 \cdot 1642$ | 670 |
| Research (collaborative or independent) | $0.20 \cdot 1694 + 0.1 \cdot 1642 + 0.05 \cdot 1642$ | 585 |
| Staff management/supervision | $0.15 \cdot 1694$ | 254 |
| Instrument availability | $1642 \cdot K-M-T$<br>K-number of instruments, M-maintenance, T- training | For capacity and billing purposes |

It is important to mention at that point that there is no immediate need to irreversibly change the job descriptions. If the situation requiring optimisation has arisen, the first step would be to review and re-allocate the duties temporarily as part of operational policy and then monitor the service. If the situation will be relieved, one can consider formal process with HR participation for updating job descriptions. If the service demand decreases (e.g. being seasonal), no changes will be required. Finally, if the demand continue to increase, the exercise will be useful in presenting administration, steering groups and funding bodies with the case to enable hiring a new member of staff or purchasing a new equipment kit. However, this example highlighted the importance of regular reviews of job descriptions according to evolving needs.

## Conclusions

A generalised mathematical approach describing the operational metrics of service (i.e. vectors for practical capacity and efficiency) presented here could be translated to any biomolecular research service, provided a matrix P (distribution of duties) is carefully gathered for each staff member of RTP and regularly reviewed to reflect changes in operational landscape. The calculation of the user access rates based on theoretical capacity is an erroneous practice and will result in substantial financial pressures for the institution and end users in the long term. In all the brevity, the user access rates for any given resource can be calculated according the very simple equation:

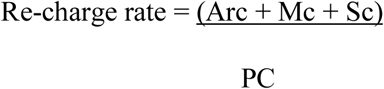

Where

Arc = Associated reagents and consumables

Mc = Maintenance contracts and repairs

Sc = Staffing costs

Pc = Practical capacity for service

(where the practical capacity is a versatile generalised metric that can be calculated according the descriptions in the appendix and excel-spreadsheet)

The general strategy how to increase performance and capacity within framework of this approach has been identified by a stepwise addressing specific components of the capacity and efficiency vectors and maximising them according to specific conditions (see supplementary figure). This strategy was illustrated with help of two specific examples: 1) under-performance due to poor operational practices and tactical execution and 2) under-performance due to the shortage of capacity caused by the outdated suboptimal operational strategy and resources.

Ultimately, this mathematical model could be adopted for enabling successful management of the entrepreneurial component of the research-business dualism manifested in the very phenomenon of core facilities and technology platforms.

## Acknowledgements

AF acknowledges support from the ISAC Emerging Leaders programme

AP acknowledges support from NIHR Cambridge Biomedical Research Centre

## Appendix. Tracking capacity and efficiency of RTP - a generalised mathematical approach

Here, a generalised mathematical approach will be presented that describes the quantitative metrics of service (i.e. capacity, utilization and efficiency) as multi-dimensional vectors that can be broken down into components and translated to benchmark, measure and monitor services provided by the biomolecular research facilities and their operational requirements (e.g. staff, equipment etc). The approach will be illustrated with practical examples. Further, it will be emphasised that the main value of this algorithm lies in the retrospective and regular monitoring of service outcomes which will allow evaluate the service in a standardised way, enumerate the requirements of the service, justify the accounting and grant applications exercises, capture the gap between theoretical and practical (not predictable) capacity, analyse the rationale behind it and address the general tactics for increasing productivity and performance in context of the existing local policies. Specific solutions will be achieved through applying this approach and identifying specific components of these vectors and maximising them according to the existing requirements. We will lay down some of the possible strategies opening a discussion how to address this issue in a systematic way, and how potentially this strategy can integrate varying operational or local procedures, and institutional or national policies.

### Definitions

For a shared laboratory resource, staffed with *n* members of staff and providing *m* different specialist services across the portfolio range, we will define:

Matrix *P*, dimension *n* × *m* for the frequency distribution of duties allocated to each particular service for each member of RTP staff *P*={*p*_*ij*_}, e.g. percentage of time dedicated by the first staff member for provision of service ‘1’, percentage of time dedicate for the provision of service ‘2’ etc:

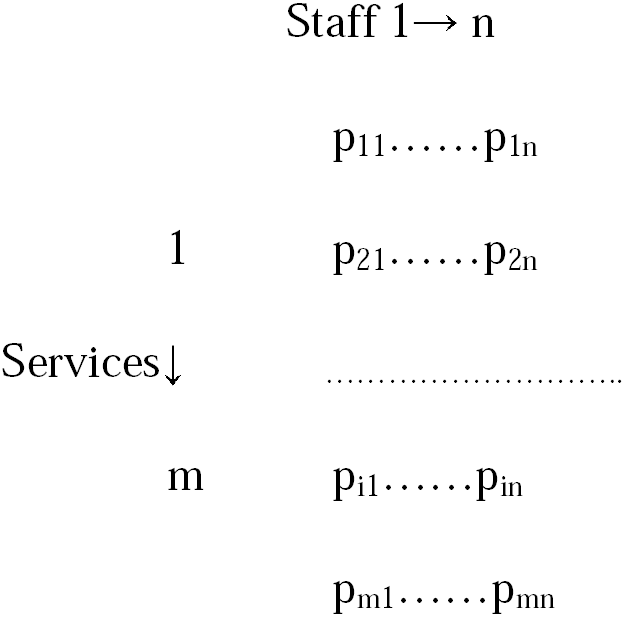

Vector *w*=*w*_*j*=_{*w*_1_……*w*_*n*_} is giving contractual annual working times (hrs) for each of *n* staff.

*w*_*a*_ averaged contractual annual working times according to local labour legislation, in hrs

*m* number of services provided

*W* number of all working days in year (365-104-public holidays)

*n* number of staff

*c* number of days RTP operations were ceased (closures due to operational reasons such as installations, maintenance, holidays, refurbishments,

*b*_*s*_ number of days the particular piece of equipment dedicated to the service was not available/malfunctioned

*k*_*s*_ number of equipment pieces dedicated to a particular service

*A*_*k*_ number of days of authorised absence for each staff member (e.g. annual leave, sickness, professional development and training etc)

*s*_*i*_= {*s*_1_……*s*_*m*_} a service output in hours over the period to be monitored/evaluated

*t*_*i*_ cumulative staff time assigned to each service, aka theoretical capacity of service

*d*_*i*_ non-bookable non-operative time for each service (e.g. contributing to ensuring that the service is running but not available to customers – for example, calibration or decontamination of a piece of equipment before the experiment)

*M*_*s*_ - time used for maintenance of a particular piece of equipment within a service s

*T*_*s*_ - time used for training on a particular piece of equipment within a service s

### Metrics of theoretical and practical capacity and efficiency

Once all the parameters have been defined (in some situations they should be already integrated into job descriptions, but obviously they can be assessed for each particular RTP/service locally through operational assessment as described in Box et al (2012) Matrix product (composition of Matrix P and vector w^t^ (transponed, row changed to column) defines the theoretical staff capacity (cumulative working time dedicated to each service for all staff)

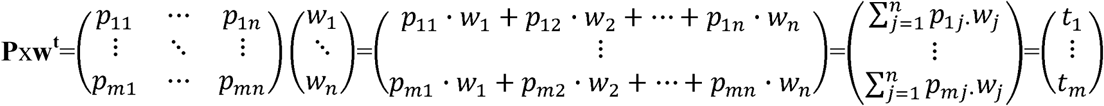

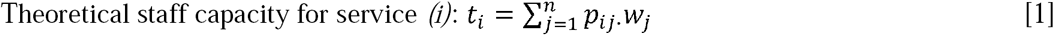

Subtracting the non-bookable non-operative time we will arrive at the cumulative theoretical cumulative operative time 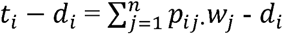

The theoretical capacity per operator per day *C* for service *(i)* can be calculated as ratio of the cumulative theoretical operative time to the product of number of staff and annual cumulative number of working days *N*=*W*·*n*

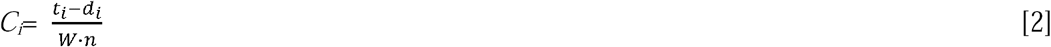

However, the practical number of operating days is different from the product of the nominal working days and number of staff (as fact, it is always due to authorised absences and other operational reasons): OD < W·n.

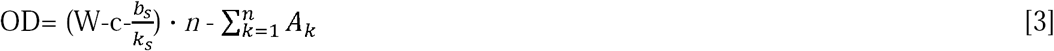

The fact of practical number of operating days being less than the number of theoretical working days requires introducing the operational efficiency as ratio of number of operational days to annual cumulative number of working days for all staff across RTP

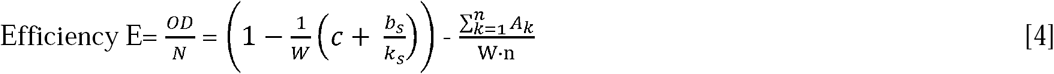

The implication of operational efficiency is that the practical staff capacity will be always less than theoretical staff capacity and needs to be defined separately as theoretical capacity per operator per day multiplied with the number of operating days OD for this particular service component.

From [4] and [5], the practical staff capacity

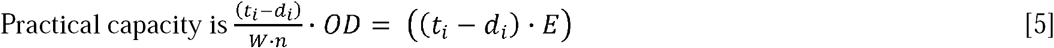

To summarise, in their vector representation theoretical and practical capacity are defined as

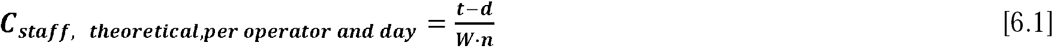

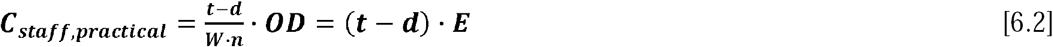

and efficiency as scalar product of OD and 1/N

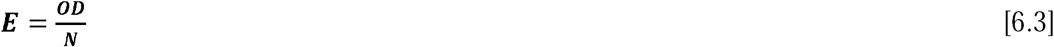

### Metrics of utilization

Utilization can be defined generally as a ratio of the service output (s in relevant units, e.g. hours, tests etc) and the staff practical capacity, and expressed in percent.

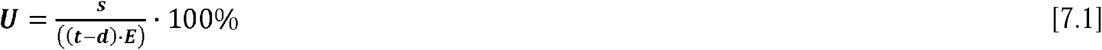

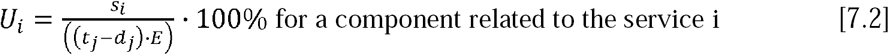

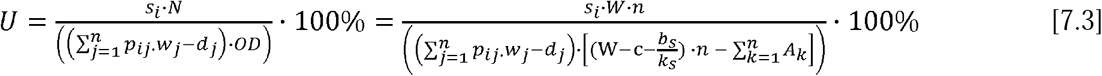

Utilization shows what fraction of the practical capacity has been realised within the review period quantified by the service output vector s.

### Review of instrumental capacity

Theoretical capacity of each instrument *C*_*instrumental, theoretical*_ within a specific service whether provided by the operator or end-user-operated is given by the product of number of the instruments *k*_*s*_ and averaged annual working contractual hours of staff *w*_*a*_:

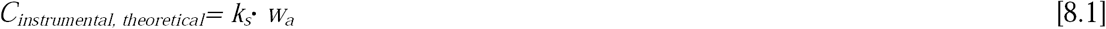

Practical capacity of each instrument *C*_*instrumental, practical*_ within a specific service whether provided by the operator or end-user-operated is given by the product of number of the instruments and averaged annual working contractual hours of staff, corrected for non-operative times such as closures and breakdowns, less training and maintenance:

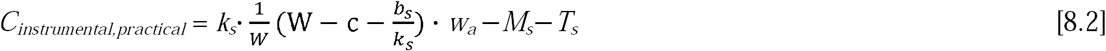

Based on the above, the rate setting for any particular service would be

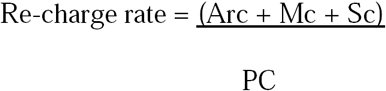

Arc = Associated reagents and consumables

Mc = Maintenance contracts and repairs

Sc = Staffing costs

Pc = Practical capacity for staff/service/instrument according the equations [6.2] and/or [8.2]

